# The long arm of childhood socioeconomic deprivation on mid- to later-life cognitive trajectories: A cross-cohort analysis

**DOI:** 10.1101/2021.06.10.447842

**Authors:** Ruby S. M. Tsang, John E. Gallacher, Sarah Bauermeister

**Author notes:** Corresponding author: Sarah Bauermeister, Department of Psychiatry, University of Oxford, Warneford Hospital, Oxford, OX3 7JX, UK.

## Abstract

**INTRODUCTION:** Earlier studies of the effects of childhood socioeconomic status (SES) on later life cognitive function consistently report a social gradient in later life cognitive function. Evidence for their effects on cognitive decline is, however, less clear.

**METHODS:** The sample consists of 5,324 participants in the Whitehall II Study, 8,572 in the Health and Retirement Study, and 1,413 in the Kame Project, who completed self-report questionnaires on their early-life experiences and underwent repeated cognitive assessments. We characterised cognitive trajectories using latent class mixed models, and explored associations between childhood SES and latent class membership using logistic regressions.

**RESULTS:** We identified distinct trajectories classes for all cognitive measures examined. Childhood socioeconomic deprivation was associated with an increased likelihood of being in a lower trajectory class.

**DISCUSSION:** Our findings support the notions that cognitive ageing is a heterogeneous process and early-life circumstances may have lasting effects on cognition across the life-course.

**Research in context:**

1. Systematic review: We reviewed the literature on childhood socioeconomic status (SES) as a predictor for cognitive decline in mid- to later-life using PubMed. Studies generally reported lower childhood SES is associated with poorer baseline cognition, but not a faster rate of decline. These studies generally focused on the mean rate of decline in the population; no study to date has explored associations between childhood SES and different cognitive trajectories. Relevant studies have been appropriately cited.
2. Interpretation: Our findings suggest that cognitive trajectories differ between individuals and across cognitive domains. Individuals of lower childhood SES were more likely to be in a lower cognitive trajectory class, which may or may not involve more rapid decline.
3. Future directions: Future studies should include more cognitive outcomes and longer follow-ups, as well as investigate the impact of social mobility to further improve our understanding on how early-life circumstances influence cognitive decline.

## 1. INTRODUCTION

Childhood adversity is known to have profound effects on cognitive development [1], with cognitive deficits or delays reported in childhood and adolescence among those who were exposed to childhood adversity [2]. Current conceptualisation of the neurodevelopmental effects of childhood adversity suggests that exposure to childhood adversity results in the dysregulation of the hypothalamic-pituitary-adrenal (HPA) axis. The HPA axis is activated and glucocorticoids are released in face of stressful experiences. With childhood adversity, the brain is exposed to prolonged periods of excessive glucocorticoids release during sensitive periods of development, which may result in lasting structural and functional changes in the brain. In addition to the HPA axis, other mechanisms such as the immune system, the microbiome as well as epigenetic alterations may also play a role in the detrimental health effects linked to child adversity [3, 4].

Given that later life cognitive function is to a great extent determined by childhood cognition [5], it has been hypothesised that the impact of childhood adversity may persist into later life, and one of the most frequently studied form of childhood adversity in ageing studies is childhood socioeconomic deprivation. Studies consistently report a social gradient in absolute later life cognitive function, with lower childhood socioeconomic status (SES) associated with poorer global cognition [6–13], memory [11, 12, 14–18], verbal fluency [17, 18], language [11], processing speed [11, 16], visuospatial abilities [11] and executive function [12] in mid- to later life. However, the literature on whether childhood SES is associated with cognitive decline is largely inconsistent. While the majority did not find an association [6, 7, 9–11, 13, 15–17, 19, 20], the few exceptions reported associations in opposite directions. For instance, one study reported that higher childhood SES was associated with slower global cognitive decline, but not with decline in specific cognitive components (episodic memory, semantic memory, and executive function) [14, 21], whereas another found that higher childhood SES was associated with faster cognitive decline [18]. Other factors such as race or sex may also modify the association. While being very poor or having poor health in childhood were not associated with faster cognitive decline, not having enough food to eat and being thinner than average in childhood were associated with slower global cognitive decline among African Americans; these effects were not observed in the Caucasians [8]. Furthermore, men in the middle childhood SES group showed faster decline in processing speed, whereas women in the low childhood SES group showed slower decline in memory and global cognition [12].

In addition, the validity of the inferences made are dependent on the validity of underlying assumptions of the models used. Studies exploring the relationships between childhood SES and cognitive decline traditionally used mixed effects or growth curve models, both of which estimate an overall mean trajectory for the entire sample and individual variation around this mean trajectory [22]. However, more recently, there is increasing evidence to suggest cognitive ageing is a heterogeneous process and distinct subgroups of trajectories exist between individuals and across cognitive domains [23, 24]. For this reason, the use of mixed effects or growth curve models may not be the most appropriate methods for modeling cognitive decline and its relationship with childhood SES.

The aim of this study is to obtain further insights into the relationship between childhood SES and cognitive decline in mid- to later life. We seek to first identify latent classes of cognitive trajectories, and then examine the predictive utility of childhood SES indicators on class membership using secondary data from three ageing cohorts, namely the Whitehall II Study [25], the Health and Retirement Study (HRS) [26] and the Kame Project [27].

## 2. METHODS

### 2.1. Cohort and study sample selection

Cohort selection was undertaken using the Dementias Platform UK (DPUK) Data Discovery tools. Our inclusion criteria were studies with:

(1) Participants aged 50 years and above;
(2) Cognitive data from three or more assessment points using the same instrument(s);
(3) Childhood SES data; and
(4) The data were already available to access on the DPUK Data Portal at the time [28].

We then investigated whether data from cohorts on other platforms may be appropriate for this study, and the relevant data were uploaded to the DPUK Data Portal with permission.

Following the aforementioned selection procedure, the Whitehall II Study, the HRS and the Kame Project were included in this study. Participants with data in at least half of the selected data collection waves were included in the analyses. Since only those aged over 65 years in HRS and over 60 years in the Kame Project completed the cognitive tests of interest, samples were restricted to those who were older than these respective age cut-offs. Participants in all three cohorts provided written informed consent at the time of data collection.

### 2.2. Cognitive outcomes

In the Whitehall II Study, cognitive data were taken from phases 7, 9, 11 and 12. Global cognition was assessed with the Mini-Mental State Examination (MMSE) [29], verbal memory with a 20-word free recall, and fluency with a 60-second written naming task of words beginning with the letter “S”. In HRS, cognitive data were taken from years 2010 to 2016. Global cognition was assessed with a modified version of the Telephone Interview for Cognitive Status (TICS-M) [30], which includes items that assess memory, attention, orientation and language, and fluency with a 60-second verbal animal naming task. In Kame, cognitive data were from the first five visits, and global cognition was assessed using the MMSE.

### 2.3. Childhood SES indicators

Participants in the three cohorts completed self-report questionnaires that included items that reflect childhood SES. The items varied between the cohorts, but covered aspects including parental education, parental unemployment and family financial hardship. The full list of these items and details on the variable coding are presented in Table A1 in the Appendix.

### 2.4. Covariates

Covariates that were tested in the models include age at the selected baseline, sex and years of education.

### 2.5. Statistical analyses

Latent class mixed models were used to identify subgroups of participants with similar cognitive performance across time. This was performed using the lcmm package version 1.7.8 [31] in R version 3.5.3. In these models, a latent class model is used to identify latent subgroups of individuals based on their trajectories and a mixed model is used to describe the mean trajectory within each subgroup simultaneously. The main underlying assumption is that the population is heterogeneous and composed of multiple latent classes with their own respective mean profiles of trajectories. These models attempt to explain the dependent variable (cognition in this case) as a function of time at the population level (fixed component), the class-specific level (mixture component), and the individual level (random component).

We used a data-driven approach adapted from methods used by Carrière et al. [32]. All models used a beta cumulative distribution function transformation to address skewness in the data. We first estimated the cognitive trajectories without adjustment for baseline covariates, then sequentially increased model complexity (intercept-only, linear or quadratic time effects for fixed, mixture and random effects, and from one to three latent classes). For each cognitive outcome, a total of 34 models were tested (see Table A2 in the Appendix). Goodness-of-fit was assessed based on model convergence, Bayesian information criterion (BIC), and average posterior probabilities (AvePP). Lower estimates of BIC indicate better model fit, and AvePP >0.7 for all trajectory classes indicate high accuracy in class assignment [33]. Then, covariates were introduced into the class-membership model in separate baseline age-adjusted models, and those with a p-value <0.20 were included in the final model. Where the final model either failed to converge or returned a smallest class being <1% of the sample, the model with the next lowest BIC value and AvePP >0.7 for all identified classes would be tested for covariates, and so forth.

After participants were classified into subgroups, logistic regressions were carried out separately with each childhood SES indicator as the predictor of class membership in Stata/SE 15.1. We accounted for multiple comparisons using the Benjamini-Hochberg procedure [34], with the false discovery rate controlled at 0.05.

All analyses were carried out in the DPUK Data Portal [28].

## 3. RESULTS

Descriptive statistics of the included participants from the three cohorts are presented in Table 1.

**Table 1.**
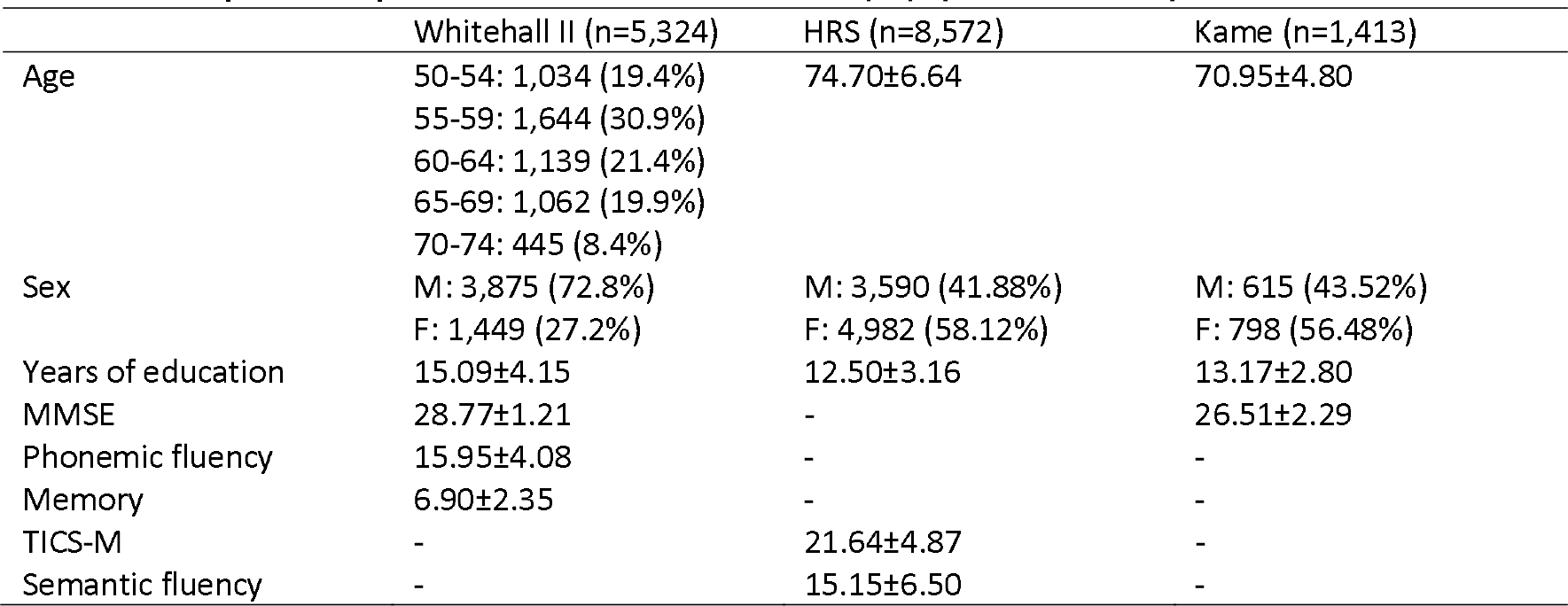
Sample descriptives at the selected baseline (n (%) or mean±s.d.).

|  | Whitehall II (n=5,324) | HRS (n=8,572) | Kame (n=1,413) |
| --- | --- | --- | --- |
| Age | 50-54: 1,034 (19.4%)<br>55-59: 1,644 (30.9%)<br>60-64: 1,139 (21.4%)<br>65-69: 1,062 (19.9%)<br>70-74: 445 (8.4%) | 74.70±6.64 | 70.95±4.80 |
| Sex | M: 3,875 (72.8%)<br>F: 1,449 (27.2%) | M: 3,590 (41.88%)<br>F: 4,982 (58.12%) | M: 615 (43.52%)<br>F: 798 (56.48%) |
| Years of education | 15.09±4.15 | 12.50±3.16 | 13.17±2.80 |
| MMSE | 28.77±1.21 | - | 26.51±2.29 |
| Phonemic fluency | 15.95±4.08 | - | - |
| Memory | 6.90±2.35 | - | - |
| TICS-M | - | 21.64±4.87 | - |
| Semantic fluency | - | 15.15±6.50 | - |

### 3.1. Characterisation of cognitive trajectories

#### 3.1.1. Whitehall II Study

In the Whitehall II Study, the best-fitting model for all three cognitive measures showed quadratic decline in the fixed and mixture components; however, they differed in the random component where there was no decline in global cognition but linear decline in fluency and memory (Table 2). Three trajectory classes were identified for global cognition and fluency, and two for memory. The patterns of trajectories appeared to be different across cognitive domains. The three global cognition classes correspond to a resilient/stable trajectory, a gradual decline trajectory, and a relatively rapid decline trajectory (Figure 1a). The three fluency classes identified represent a resilient/stable trajectory, a gradual decline trajectory, and a curvilinear trajectory showing an initial improvement followed by rapid decline (Figure 1b). The two memory classes identified both showed decline over time (Figure 1c).

**Table 2.**
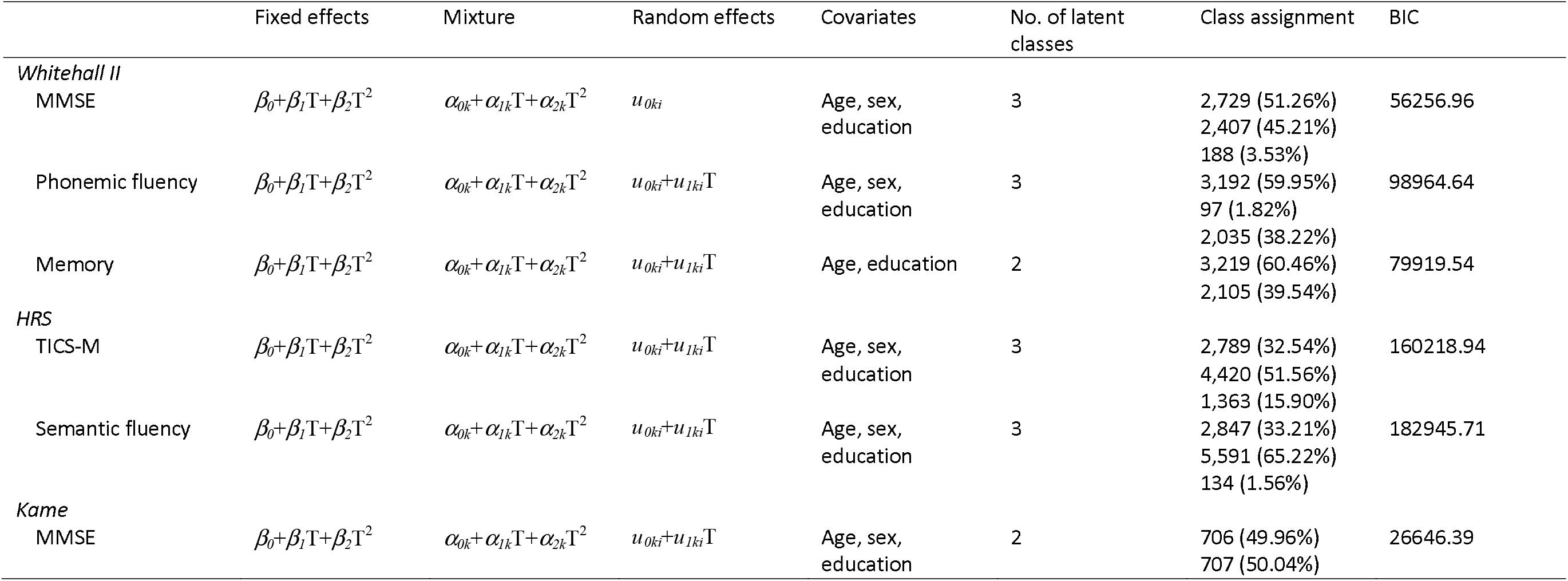
Parameters, model fit indices and class assignment in the final LCMM models.

**Figure 1.**
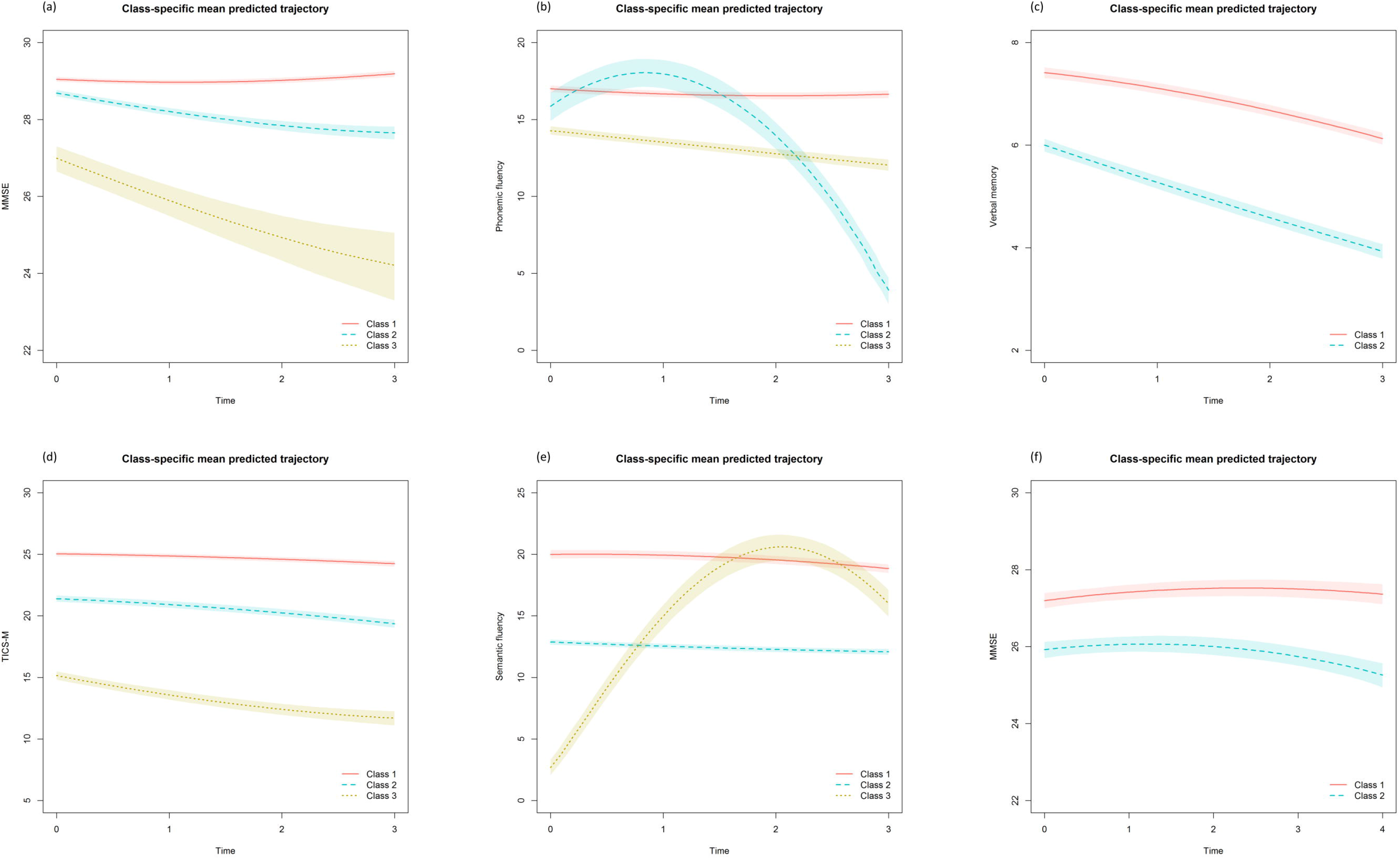
Mean predicted trajectories for the identified classes in Whitehall II: (a) MMSE, (b) phonemic fluency, (c) memory; in HRS: (d) TICS-M, (e) semantic fluency; and in Kame: (f) MMSE.

#### 3.1.2. Health and Retirement Study

The best-fitting model for both global cognition and fluency in HRS showed quadratic decline in the fixed and mixture components, and linear decline in the random component (Table 2). Both models identified three trajectory classes. The three global cognition classes all showed gradual decline (Figure 1d), whereas two of the three fluency classes showed gradual decline, and the third showed some initial improvement followed by rapid decline (Figure 1e).

#### 3.1.3. Kame Project

Similar to most of the other cognitive measures examined, the best-fitting model for global cognition in the Kame Project showed quadratic decline in the fixed and mixture components, and linear decline in the random component. The two trajectory classes correspond to a resilient/stable trajectory and a gradual decline trajectory (Figure 1f).

### 3.2. Associations between exposure to early adversity and class membership

Using Class 1 as the reference group, it appeared that among the childhood SES indicators examined, almost all showed an association between lower childhood SES and increased likelihood of membership in a lower trajectory class (Table 3).

**Table 3.** Associations between childhood SES indicators and the likelihood of being in a lower trajectory (with the top trajectory ‘Class 1’ as reference)

| Childhood SES indicator | Global cognition<br>Class 2 |  | Global cognition<br>Class 3 |  | Fluency Class 2 |  | Fluency Class 3 |  | Memory Class 2 |  |
| --- | --- | --- | --- | --- | --- | --- | --- | --- | --- | --- |
|  | *B* | *p* | *B* | *p* | *B* | *p* | *B* | *p* | *B* | *p* |
| *Whitehall II* |  |  |  |  |  |  |  |  |  |  |
| Age when father finished full-time education | -0.06* | <0.0001 | -0.05 | 0.2392 | -0.03 | 0.5568 | -0.10* | <0.0001 | -0.08* | <0.0001 |
| Age when mother finished full-time education | -0.11* | <0.0001 | -0.26* | 0.0001 | -0.11 | 0.1057 | -0.18* | <0.0001 | -0.14* | <0.0001 |
| Father's social class | -0.11* | 0.0001 | -0.24* | 0.0017 | -0.12 | 0.2589 | -0.16* | <0.0001 | -0.10* | 0.0003 |
| Spent four or more weeks in hospital | 0.14 | 0.1126 | 0.13 | 0.5800 | 0.46 | 0.1098 | 0.38* | <0.0001 | 0.35* | <0.0001 |
| Father/mother were unemployed when they wanted to be working | 0.31* | 0.0010 | 0.59* | 0.0082 | -0.68 | 0.1420 | 0.13 | 0.1629 | 0.27* | 0.0031 |
| Family had continuing financial problems | 0.17* | 0.0073 | 0.36* | 0.0309 | 0.15 | 0.5305 | 0.29* | <0.0001 | 0.31* | <0.0001 |
| Family/household did not have an inside toilet | 0.36* | <0.0001 | 0.74* | <0.0001 | 0.40 | 0.0973 | 0.50* | <0.0001 | 0.43* | <0.0001 |
| Family/household owned a car | -0.60* | <0.0001 | -1.28* | <0.0001 | -0.65* | 0.0028 | -0.97* | <0.0001 | -0.95* | <0.0001 |
| *HRS* |  |  |  |  |  |  |  |  |  |  |
| Father's education | -0.12* | <0.0001 | -0.27* | <0.0001 | -0.14* | <0.0001 | -0.07 | 0.1023 |  |  |
| Mother's education | -0.14* | <0.0001 | -0.32* | <0.0001 | -0.16* | <0.0001 | -0.14* | 0.0009 |  |  |
| Childhood health | -0.20* | <0.0001 | -0.45* | <0.0001 | -0.24* | <0.0001 | -0.08 | 0.3850 |  |  |
| Family financially poor | 0.50* | <0.0001 | 0.82* | <0.0001 | 0.48* | <0.0001 | 0.32 | 0.0881 |  |  |
| Family moved due to financial difficulties | 0.30* | <0.0001 | 0.33* | 0.0001 | 0.14* | 0.0210 | 0.29 | 0.1856 |  |  |
| Family received help because of financial difficulties | -0.01 | 0.8708 | -0.19 | 0.0745 | -0.20* | 0.0036 | -0.07 | 0.7791 |  |  |
| Father unemployed | 0.32* | <0.0001 | 0.37* | <0.0001 | 0.28* | <0.0001 | 0.25 | 0.2458 |  |  |
| *Kame* |  |  |  |  |  |  |  |  |  |  |
| Father's education | -0.09* | <0.0001 |  |  |  |  |  |  |  |  |
| Mother's education | -0.10* | <0.0001 |  |  |  |  |  |  |  |  |
| Household density | 0.07 | 0.1734 |  |  |  |  |  |  |  |  |
| Urban/suburban living | -0.41* | 0.0003 |  |  |  |  |  |  |  |  |
| Family financial difficulties | 0.04 | 0.0556 |  |  |  |  |  |  |  |  |

\* *FDR* < 0.05

In the Whitehall II Study, older age when father completed full-time education, older age when mother completed full-time education, higher father’s social class and family car ownership were consistently associated with a decreased likelihood of being in a lower trajectory class across cognitive domains, while ongoing family financial problems, and not having an inside toilet in the household were associated with an increased likelihood of being in a lower trajectory class. Having spent four or more weeks in hospital and parental unemployment were associated with a higher likelihood of being in a lower trajectory class in two out of three cognitive outcomes. There was limited evidence for an association between childhood SES and class membership in the small trajectory class with initial learning effects followed by rapid decline in the fluency task.

Similarly, almost all childhood SES indicators examined were associated with an increased likelihood of membership in a lower trajectory class in the HRS. Higher father’s education, higher mother’s education, better childhood health were consistently associated with a decreased likelihood of being in a lower trajectory class. Worse family financial status, family having moved due to financial difficulties, and father’s unemployment were related to an increased likelihood of being in a lower trajectory class. Again, there was limited evidence for an association between childhood SES and class membership in the trajectory class with initial learning effects followed by rapid decline in the fluency task.

The results from Kame, however, present a mixed picture. On one hand, higher father’s education, higher mother’s education, and urban/suburban living were linked to a lower likelihood of being in the lower trajectory class. On the other hand, household density and family financial situation were not associated with the likelihood of being in the lower trajectory class.

## 4. DISCUSSION

The primary aim of this study is to explore the relationship between childhood SES and mid- to late-life cognitive trajectories. Using longitudinal data from three cohorts, we characterised latent cognitive trajectories, and examined associations between childhood SES indicators and cognitive trajectory class membership. We found: (i) there were multiple trajectory classes in all of the cognitive outcomes included in this study, and (ii) lower childhood SES is consistently associated with an increased likelihood of being in a lower trajectory class.

Earlier studies that examined cognitive ageing or cognitive decline typically used statistical methods that assume there is one mean trajectory within the population. In contrast to these studies, we found that there are multiple latent trajectory classes, but the profiles observed differed between cohorts and across cognitive domains. This provides further support for the notion that cognitive ageing is a heterogeneous process, with between- and within-cohort variation. It is important to note that for certain cognitive outcomes (e.g., memory in the Whitehall II Study and TICS-M in HRS), the slopes of the different predicted trajectories appeared rather similar, suggesting that rates of cognitive decline may not vary substantially.

An interesting finding in the patterns of cognitive trajectories observed, was that in the fluency tasks in both the Whitehall II Study and HRS, there was a small class that showed a curvilinear trajectory showing initial improvements followed by more rapid decline. These initial improvements may reflect practice effects from repeated administration of cognitive assessments. Studies have found considerable practice effects even when assessments were conducted several years apart, and such short-term improvements are often large enough to counteract age-related cognitive decline [35]. While it may seem counterintuitive that those who will eventually be cognitively impaired in fact show greater practice effects, one explanation is that these individuals were performing below their actual cognitive potential when they first encounter novel cognitive tests, as they need more time to understand the task demands. As they familiarise themselves with the task characteristics, they then exhibit a “rebound” in their performance (i.e., a “novelty effect) [36, 37]. Thorgusen and colleagues [38] demonstrated that both memory and novelty effects uniquely contribute towards these neuropsychological practice effects, and cognitive impairment is more likely to be associated with smaller practice effects in memory tasks, but larger practice effects in tasks assessing other cognitive domains. It has been proposed that such novelty effect may be a useful early marker of declining cognitive reserve and neurodegeneration, but more research is required to understand how practice effects differ depending on the population, task complexity, and cognitive domain assessed before conclusions can be drawn about their potential diagnostic and prognostic utility.

Our results suggest that childhood SES is an important contributor to mid- to later-life cognitive trajectories. Level of education is often used as a measure of early-life SES; there is consistent evidence that education plays an important role on later-life cognitive function and cognitive decline [39–41], but few studies have examined the effects of other measures of childhood SES, especially on cognitive decline. This paper adds to the literature by including a range of childhood SES indicators, and examining their associations with cognitive trajectories. Major strengths of this study also include larger sample sizes than earlier studies, and the cross-cohort comparisons also showed that the associations between childhood SES indicators and different cognitive trajectory classes were robust across cohorts and cognitive domains. The finding that there are distinct latent trajectory classes that may not necessarily differ in their slopes also help explain some of the inconsistencies in earlier studies that while most studies reported no association between childhood SES and rate of cognitive decline [6, 7, 9–11, 13, 15–17, 19, 20], a few others found associations in opposite directions [14, 18, 21].

These findings have implications for the prevention of cognitive impairment and dementia. Globally, the number of people living with dementia is rapidly increasing, but there is currently no cure and no treatment that alters the course of the disease. Delaying or preventing the onset of dementia is therefore a key public health priority. Our analyses showed that the main difference between latent trajectory classes generally lies in their baseline cognitive function, and lower childhood SES is consistently associated with the lower trajectory classes. This suggests that childhood SES is an important contributor to cognitive reserve, and interventions aimed at reducing socioeconomic inequalities may be effective in delaying or preventing the onset of dementia. Furthermore, studies using a life-course approach have demonstrated that SES at different life stages each make unique contributions to cognitive function in mid- to later-life, and upward social mobility later in life may to a certain extent counteract the negative effects of disadvantaged childhood SES [42, 43]. Whether upward social mobility later in life influences mid- to later-life cognitive decline remains to be investigated.

Finally, some theoretical and methodological issues should be addressed. First, observable cognitive change across time is partly dependent on the psychometric properties of the cognitive instruments used. For instance, MMSE is known to have a strong ceiling effect [44] and shows poor sensitivity to change in the tails of the distribution [45, 46]. The curvilinear nature of the instrument means that a one-point change in the higher range of scores do not hold the same clinical meaning as a one-point change in the medium or lower range, it is possible that cognitive scores may need to be appropriately transformed before a fair comparison can be made between the slopes of modelled cognitive trajectories.

Second, the magnitude of cognitive change observed is somewhat dependent on the length and frequency of follow-up, as well as the demographic characteristics of the sample. We modelled cognitive decline over four waves of data in both the Whitehall II Study and HRS, and five waves in the Kame Project. However, this corresponds to around 12, six and eight years in the respective studies. Such difference in follow-up durations likely explains the more evident cognitive decline observed in the Whitehall II Study compared to the other two cohorts. Moreover, the three cohorts exhibit large demographic differences between them, while the Whitehall II Study is a cohort of British civil servants, the HRS is a longitudinal panel study that surveyed a representative sample in the United States, and the Kame Project is a cohort of older Japanese Americans in the United States. These sampling differences have resulted in differences in the age and sex distributions within the cohorts, as well as differences in the level of education of the participants, which may all have effects on cognitive ageing trajectories.

In summary, different patterns of cognitive decline were observed between cohorts and across cognitive domains, and lower childhood SES generally predicted membership in a lower cognitive trajectory class. Future research may benefit from examining trajectories using different cognitive instruments with better psychometric properties as well as assess more cognitive domains than what we have examined here, including data from longer follow-ups, and exploring the influence of social mobility on mid- to later-life cognitive trajectories.

## Supporting information

Appendix Tsang

Highlights Tsang

## Acknowledgements

Funding: RT was supported by Dementias Platform UK (DPUK) and all analyses were conducted on the DPUK Data Portal. This study was conducted under DPUK Study 0144 as part of the Early Adversity and Dementia Research Programme. The Medical Research Council supports DPUK through grant MR/L023784/2.

