## Appendix Tsang for "The long arm of childhood socioeconomic deprivation on mid- to later-life cognitive trajectories: A cross-cohort analysis"

**Table A1. Childhood SES indicators in Whitehall II, HRS and Kame.**

| Cohort | Item | Coding |
| --- | --- | --- |
| Whitehall II | How old was your father when he finished full-time education? | (Top-coded; range=11-25) |
|  | How old was your mother when she finished full-time education? |  |
|  | What is/was your father’s main job, what kind of work does/did he do in it? (Occupations classified into Registrar-General’s social classes.) | 1=V  2=IV  3=III manual  4=III non-manual  5=II  6=I |
|  | You spent four or more weeks in hospital. | 0=No  1=Yes |
|  | Your father/mother were unemployed when they wanted to be working. |  |
|  | Your family had continuing financial problems. |  |
|  | Your family/household did not have an inside toilet. |  |
|  | Your family/household owned a car. |  |
| HRS | What is the highest grade of school your father completed? | 0=No formal education  1-11=Grades  12=High school  13-15=Some college  16=College grad  17=Post-college |
|  | And what is the highest grade of school your mother completed? |  |
|  | Consider your health while you were growing up, from birth to age 16. Would you say that your health during that time was excellent, very good, good, fair, or poor? | 1=Poor  2=Fair  3=Good  4=Very good  5=Excellent |
|  | Now think about your family when you were growing up, from birth to age 16. Would you say your family during that time was pretty well off financially, about average, or poor? | 0=Pretty well off financially or about average  1=Poor |
|  | While you were growing up, before age 16, did financial difficulties ever cause you or your family to move to a different place? | 0=No  1=Yes |
|  | Before age 16, was there a time when you or your family received help from relatives because of financial difficulties? |  |
|  | Before age 16, was there a time of several months or more when your father had no job?^a^ |  |
| Kame | What was the highest grade or year of regular school your father completed? | 1-8=Elementary, first to eighth grade  9-12=High school, first to fourth year  13-16=College, first to fourth year  19-21=Doctorate, first to third year |
|  | What was the highest grade or year of regular school your mother completed? | 1-8=Elementary, first to eighth grade  9-12=High school, first to fourth year  13-16=College, first to fourth year  19-21=Doctorate, first to third year |
|  | Including yourself, how many people lived in your house when you were 2 to 3 years old?^b^ | Top-coded: continuous until 11 = >10 |
|  | How many bedrooms were in this house?^b^ | Top-coded: continuous until 8 = >7 |
|  | During most of your growing up years, did you live on a farm, in a rural area, small town, or large city? | 0 = On a farm with animals (cattle, horses, pigs, etc.) or in a rural area (e.g. an orchard or berry farm, or non-farm)  1 = In a small town (or suburb) (<10,000) or in a large city |
|  | On a scale from 1 to 10, can you rate your family’s financial situation when you were a child, with 1 being no financial difficulty and 10 being extremely difficult? |  |

^a^ “Father never worked/always disabled” was recoded as 1 and “Never lived with father/father was not alive” was recoded as missing.

^b^ A household density variable was derived using these two variables.

**Table A2. LCMM models tested.**

| Fixed effects | Mixture (only applicable when ≥2 classes) | Random effects | Number of latent classes | Covariates tested (individually then combined) |
| --- | --- | --- | --- | --- |
| *β_0_* | *α_0k_* | *u_0ki_* | 1-3 | Age, sex, years of education |
| *β_0_*+*β_1_*T | *α_0k_* | *u_0ki_* | 1-3 | Age, sex, years of education |
| *β_0_*+*β_1_*T | *α_0k_* | *u_0ki_*+*u_1ki_*T | 1-3 | Age, sex, years of education |
| *β_0_*+*β_1_*T | *α_0k_*+*α_1k_*T | *u_0ki_* | 1-3 | Age, sex, years of education |
| *β_0_*+*β_1_*T | *α_0k_*+*α_1k_*T | *u_0ki_*+*u_1ki_*T | 1-3 | Age, sex, years of education |
| *β_0_*+*β_1_*T+*β_2_*T^2^ | *α_0k_* | *u_0ki_* | 1-3 | Age, sex, years of education |
| *β_0_*+*β_1_*T+*β_2_*T^2^ | *α_0k_* | *u_0ki_*+*u_1ki_*T | 1-3 | Age, sex, years of education |
| *β_0_*+*β_1_*T+*β_2_*T^2^ | *α_0k_* | *u_0ki_*+*u_1ki_*T+*u_2ki_*T^2^ | 1-3 | Age, sex, years of education |
| *β_0_*+*β_1_*T+*β_2_*T^2^ | *α_0k_*+*α_1k_*T | *u_0ki_* | 1-3 | Age, sex, years of education |
| *β_0_*+*β_1_*T+*β_2_*T^2^ | *α_0k_*+*α_1k_*T | *u_0ki_*+*u_1ki_*T | 1-3 | Age, sex, years of education |
| *β_0_*+*β_1_*T+*β_2_*T^2^ | *α_0k_*+*α_1k_*T | *u_0ki_*+*u_1ki_*T+*u_2ki_*T^2^ | 1-3 | Age, sex, years of education |
| *β_0_*+*β_1_*T+*β_2_*T^2^ | *α_0k_*+*α_1k_*T+*α_2k_*T^2^ | *u_0ki_* | 1-3 | Age, sex, years of education |
| *β_0_*+*β_1_*T+*β_2_*T^2^ | *α_0k_*+*α_1k_*T+*α_2k_*T^2^ | *u_0ki_*+*u_1ki_*T | 1-3 | Age, sex, years of education |
| *β_0_*+*β_1_*T+*β_2_*T^2^ | *α_0k_*+*α_1k_*T+*α_2k_*T^2^ | *u_0ki_*+*u_1ki_*T+*u_2ki_*T^2^ | 1-3 | Age, sex, years of education |
