## Supplementary material for "The long arm of childhood socioeconomic deprivation on mid- to later-life cognitive trajectories: A cross-cohort analysis": Highlights Tsang

- Studies consistently report a social gradient in later life cognitive function.
- Effects of childhood socioeconomic deprivation on cognitive decline is less clear.
- We identified distinct cognitive trajectories for all cognitive domains examined.
- Those from a deprived childhood were more likely to be in a lower trajectory class.
